# Caffeine stabilises fission yeast Wee1 in a Rad24-dependent manner but attenuates its expression in response to DNA damage identifying a putative role for TORC1 in mediating its effects on cell cycle progression

**DOI:** 10.1101/2020.01.22.915231

**Authors:** John P. Alao, Johanna Johansson-Sjölander, Charalampos Rallis, Per Sunnerhagen

## Abstract

The widely consumed neuroactive compound caffeine has generated much interest due to its ability to override the DNA damage and replication checkpoints. Previously Rad3 and its homologues was thought to be the target of caffeine’s inhibitory activity. Later findings indicate that the Target of Rapamycin Complex 1 (TORC1) is the preferred target of caffeine. Effective Cdc2 inhibition requires both the activation of the Wee1 kinase and inhibition of the Cdc25 phosphatase. The TORC1, DNA damage, and environmental stress response pathways all converge on Cdc25 and Wee1. We previously demonstrated that caffeine overrides DNA damage checkpoints by modulating Cdc25 stability. The effect of caffeine on cell cycle progression resembles that of TORC1 inhibition. Furthermore, caffeine activates the Sty1 regulated environmental stress response. Caffeine may thus modulate multiple signalling pathways that regulate Cdc25 and Wee1 levels, localisation and activity. Here we show that the activity of caffeine stabilises both Cdc25 and Wee1. The stabilising effect of caffeine and genotoxic agents on Wee1 was dependent on the Rad24 chaperone. Interestingly, caffeine inhibited the accumulation of Wee1 in response to DNA damage. Caffeine therefore modulates cell cycle progression contextually through increased Cdc25 activity and Wee1 repression following DNA damage via TORC1 inhibition.

## Introduction

Cell cycle progression through mitosis is under the opposing control of the Cdc25 phosphatase and the Wee1 kinase. Cdc25 removes inhibitory phosphorylation moieties on Cdc2, which in turn enhances Cdc25 activity in a positive feedback loop. In contrast, Wee1 phosphorylates Cdc2 on tyrosine residue 15 to inhibit its activity. Cdc2 in turn negatively regulates Wee1 by phosphorylation leading to its nuclear exclusion or degradation (Caspari & Hilditch, 2015, Moseley, 2017, de Gooijer *et al*., 2017). Cells must delay progress through S-phase and mitosis in response to stalled replication, DNA double strand breaks and other forms of damage, in order to effect DNA repair and maintain viability (Karlsson-Rosenthal & Millar, 2006, Alao & Sunnerhagen, 2008). Effective activation and maintenance of DNA damage checkpoints thus involves the dual regulation of both Cdc25 and Wee1 via a “double lock” mechanism (Raleigh & O’Connell, 2000). Activation of the DNA damage response pathway induces inhibitory Cdc25 phosphorylation, Rad24 binding, nuclear export and stockpiling within the cytoplasm. In contrast, increased Wee1 activation occurs via phosphorylation, and this kinase accumulates within the nucleus (Karlsson-Rosenthal & Millar, 2006, Alao & Sunnerhagen, 2008). Caffeine has generated much controversy by its ability to override checkpoint signalling but the underlying mechanisms remain unclear. Caffeine inhibits members of the members of the phosphatidylinositol 3 kinase-like kinase (PIKK) family including ataxia telangiectasia mutated (ATM) and ataxia – and rad related (ATR) kinase homologue Rad3 and Target of Rapamycin Complex 1 (TORC1) *in vitro* (Humphrey, 2000, Bode & Dong, 2007, Lovejoy & Cortez, 2009, Gibbs *et al*., 2015). Initial reports suggested that caffeine overrides DNA damage checkpoint signalling by inhibiting *Schizosaccharomyces pombe* Rad3 and its orthologues but this view remains controversial (Moser *et al*., 2000, Wanke *et al*., 2008, Cortez, 2003). Studies that are more recent indicate that TORC1 is the major cellular target of caffeine *in vivo* (Kuranda *et al*., 2006, Reinke *et al*., 2006, Wanke *et al*., 2008, Rallis *et al*., 2013). TORC1 is a major regulator of cell cycle progression acting on both Cdc25 and Wee1. The inhibition of TORC1 activity suppresses Wee1 expression, results in increased Cdc25 activation and drives cells into mitosis. In addition, the effect of caffeine on cell cycle progression resembles that of TORC1 inhibition (Petersen, 2009, Alao *et al*., 2014, Atkin *et al*., 2014). We previously demonstrated that caffeine overrides checkpoint signalling in part, by stabilising Cdc25 expression. Interestingly, deletion of the *rad3*^+^ gene and its downstream target Cds1 similarly resulted in Cdc25 stabilisation. These findings suggested a role of Rad3 signalling in regulating Cdc25 stability during the normal cell cycle. Similarly, the Sty1-regulated Environmental Stress Response (ESR) pathway also plays a role in regulating both Cdc25 and Wee1 expression levels and is activated by caffeine (Alao & Sunnerhagen, 2008, Paul *et al*., 2018). The integration of Cdc25 and Wee1 phosphorylation, localisation, stability and activity thus play a key role in modulating the timing of mitosis. TORC1, Rad3 and Sty1 regulate the major signalling pathways that converge on the Cdc25 and Wee1 axis (Alao & Sunnerhagen, 2008, Petersen, 2009, de Gooijer *et al*., 2017). Herein we further explored the mechanism(s) by which caffeine stabilises Cdc25 and overrides checkpoint signalling, and have investigated the impact of subcellular localisation under these conditions. Here we report that in addition to modulating Cdc25 activity, caffeine also suppressed the DNA damage-induced stabilisation of Wee1 in *S. pombe*. In contrast, caffeine stabilised Wee1 in a Rad24 dependent manner under normal cell cycle conditions. These findings demonstrate that caffeine overrides the DNA damage checkpoints by positively regulating Cdc25 and negatively regulating Wee1. They also provide further evidence for the assertion that caffeine modulates TORC1 (and other pathways) and not Rad3 signalling to overcome the DNA damage checkpoint “double lock” mechanism.

## Results

### Caffeine stabilises Cdc25 by inhibiting its nuclear degradation

We previously demonstrated that caffeine stabilises both wild type (wt) Cdc25-GFP_int_ and the Cdc25_(9A)_-GFP_int_ mutant that lacks 9 major inhibitory phosphorylation sites and is normally degraded following exposure to genotoxic agents (Frazer & Young, 2011, Frazer & Young, 2012, Alao *et al*., 2014). In this study, exposure to 10 mM caffeine also stabilised the Cdc25_(12A)_ mutant protein that lacks all 12 inhibitory phosphorylation sites (Fig. 1 A). Caffeine also stabilised Cdc25 in *mik1Δ* mutants. Mik1 is required for maintenance of the replication damage checkpoint signalling in *S. pombe* mutants expressing Cdc25_(12A)_-GFP_int_ (Frazer & Young, 2011, Frazer & Young, 2012). As observed for *rad3*Δ and *cds1*Δ mutants (Alao *et al*., 2014), Cdc25 appeared more stable in a *mik1*Δ genetic background (Fig. 1 A). Caffeine also suppressed the 20 mM hydroxyurea (HU)-induced degradation of the Cdc25_(12A)_-GFP_int_ mutant (Fig. 1 B). The stabilising effect of caffeine was not due to stress-induced Sty1 activation, as exposure to 0.6 M KCl induced Cdc25_(9A)_-GFP_int_ degradation (Fig. 1 C). Inhibition of Crm1-dependent nuclear export with 100 ng/ml leptomycin B (LMB) slightly suppressed Cdc25 levels in both wt and Cdc25_(12A)_-GFP_int_ expressing mutants (Fig. 1 D). LMB also inhibited the stockpiling of Cdc25-GFP_int_ following exposure to HU but failed to stabilise Cdc25_(12A)_-GFP_int_ under these conditions (Fig. 1 D). Similarly, HU-induced stockpiling of Cdc25 was dependent on Rad24 (Fig. 1 E). As previously reported, caffeine is more effective at overriding DNA damage checkpoints in strains expressing Cdc25 mutant protein that cannot be negatively phosphorylated (Fig. 1 F,G).

**Figure 1.**
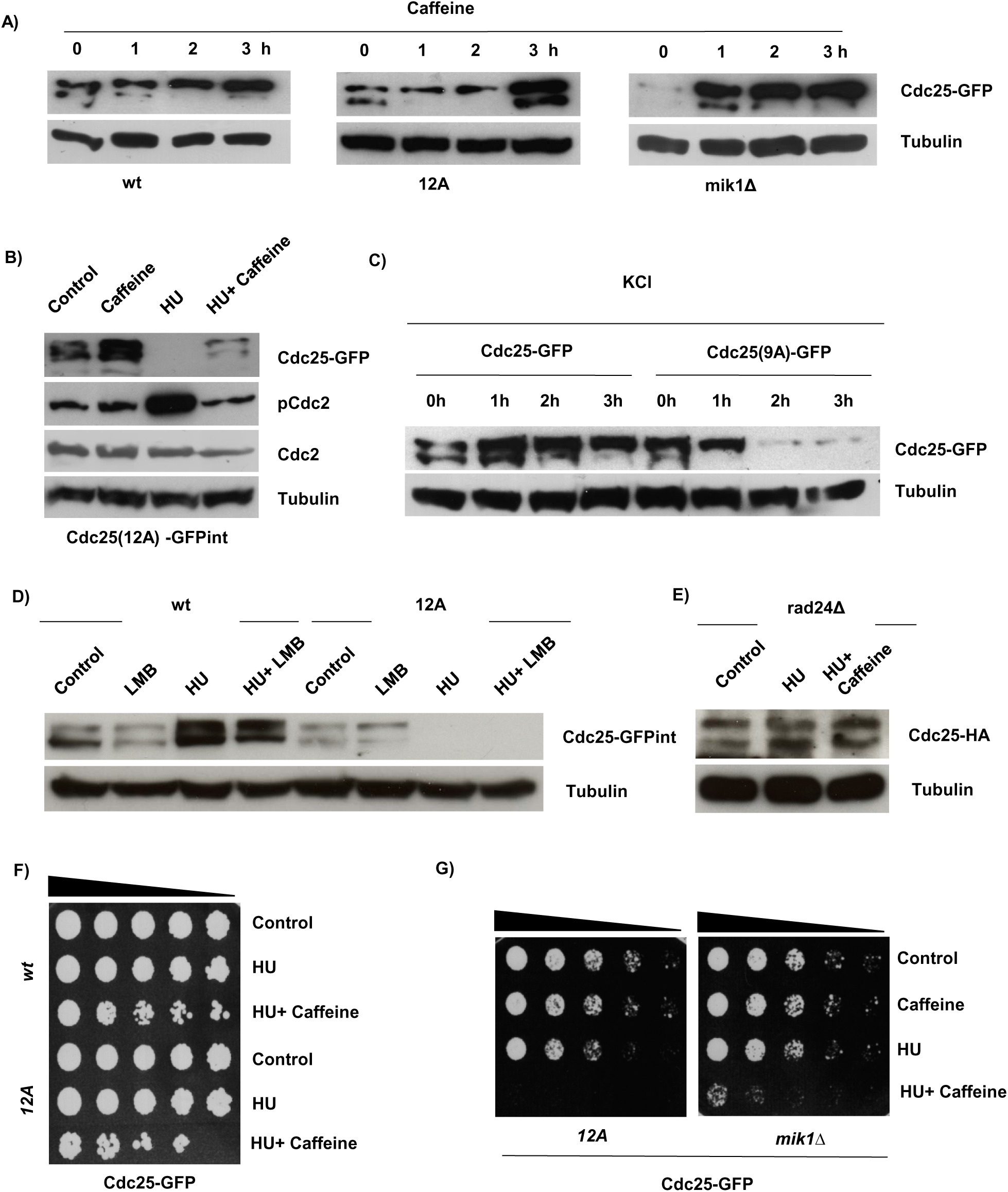
Caffeine induces the nuclear accumulation of Cdc25 in *S. pombe*. **A.** Strains expressing wt Cdc25-GFP, Cdc25_(12A)_-GFP, or Cdc25-GFP from a *mik1*Δ genetic background were incubated with 10 mM caffeine and harvested at the indicated time points. Total protein lysates were resolved by SDS-PAGE and Cdc25 detected using antibodies directed against GFP. Gel loading was monitored using antibodies directed against tubulin. **B.** Strains expressing Cdc25_(12A)_-GFP_int_ were pre-treated with 20 mM HU for two hours, followed by the addition of 10 mM caffeine. Cultures were incubated for a further 2 hours and total protein lysates were resolved by SDS-PAGE. Cdc25, phospho-Cdc2 and Cdc2 were detected using antibodies directed against GFP. Gel loading was monitored as in **A**. **C.** Strains expressing wt Cdc25-GFP or Cdc25_(9A)_-GFP were cultured in YES containing 0.6 M KCl and harvested at the indicated time points. Samples were analysed as in **A**. **D.** Strains expressing wt Cdc25-GFP or Cdc25_(12A)_-GFP_int_ were incubated with 100 ng/ml LMB and 20 mM HU alone or in combination. Cells were pre-treated with HU for 2 h and then incubated with LMB for another 2 h as indicated. Samples were processed as in **A**. **E.** The Cdc25-HA *rad24*Δ strain was exposed to 20 mM HU alone or in combination with 10 mM caffeine. Cells were pre-treated with HU for 2 h and then incubated with caffeine for another 2 h as indicated. Cdc25 was detected using antibodies directed against the HA epitope. Gel loading was monitored using antibodies directed against tubulin. **F.** Strains expressing wt Cdc25-GFP and Cdc25_(12A)_-GFP_int_ exposed to 20 mM HU alone or in combination with 10 mM caffeine as in **E**. Samples were adjusted for relative cell numbers, serially diluted, plated on YES agar and incubated for 2-3 days. **G.** Cdc25_(12A)_-GFP_int_ and Cdc25-GFP expressing strains from a *mik1*Δ genetic background were treated as in **F**.

### Caffeine stabilises Wee1 in a Rad24 dependent manner

TORC1 inhibition activates Cdc25 and suppresses Wee1 activity (Atkin *et al*., 2014). As Cdc25 and Wee1 are both partially regulated by ubiquitin-dependent degradation, we next investigated the effect of caffeine on Wee1 expression. Exposure to caffeine induced a rapid and time-dependent increase in Wee1 levels. Wee1 levels increased with 30 min of exposure to caffeine and continued to rise over the course of the 4 h incubation periods (Fig. 2 A, B). The caffeine-induced accumulation of Wee1 was dependent on Rad24 expression (Fig. 2 C). A role for Rad24 in regulating Wee1 stability has previously been reported (Paul *et al*., 2018). Caffeine may also affect the phosphorylation of Wee1 in a manner that is distinct from a general effect on protein degradation (Lucena *et al*., 2017). The deletion of *rad24^+^* resulted in a partial reduction of Wee1 expression and induced a “wee” phenotype (Fig. 2 D, E). Additionally, inhibition of nuclear export with 100 ng/ml LMB stabilised Wee1 independently of Rad24 (Fig. 2 F). Caffeine also stabilised Mik1 under normal cell cycle conditions (Fig. 2 G). Caffeine thus interferes with the coordinated regulation of Cdc25 and Wee1, possibly via inhibition of TORC1 activity.

**Figure 2.**
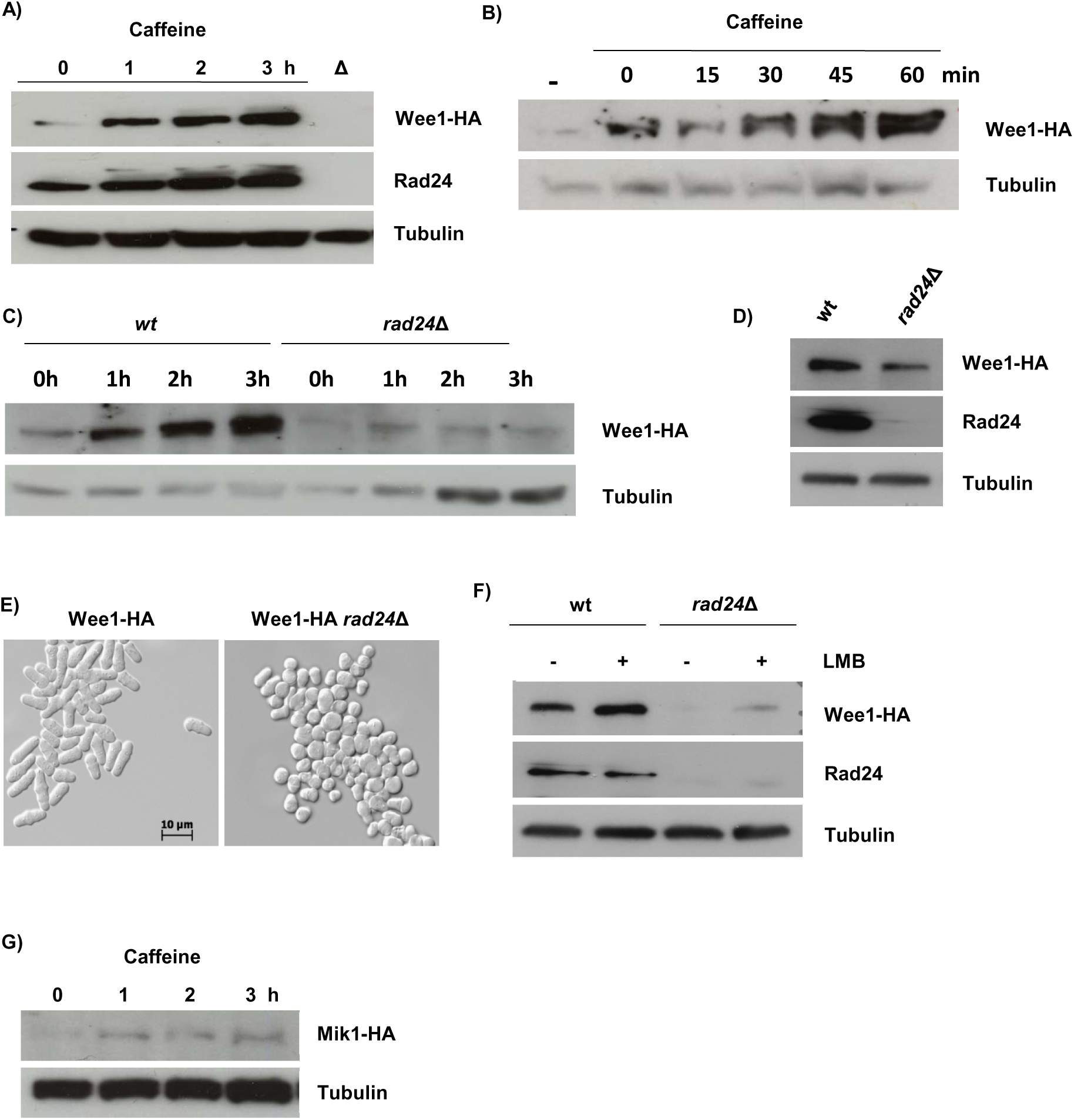
Caffeine induces Wee1 accumulation in a Rad24-dependent manner in *S. pombe*. **A.** A Wee1-HA expressing strain was incubated with 10 mM caffeine and harvested at the indicated time points. Total protein lysates were resolved by SDS-PAGE. Wee1 was detected using antibodies directed against the HA epitope. Rad24 was detected using a pan 14-3-3 antibody. A *rad24*Δ mutant was used to monitor antibody specificity. Gel loading was monitored using antibodies directed against tubulin. **B.** Cells expressing Wee1-HA were treated as in **A**. **C.** Wt and *rad24*Δ mutant cells expressing Wee1-HA were treated as in **A**. **D.** Wt and *rad24*Δ cells expressing Wee1-HA were grown to log phase and treated as in **A**. **E.** Log phase cultures of wt and *rad24*Δ cells were fixed in ethanol and examined by differential contrast microscopy. **F.** Wt and *rad24*Δ strains expressing Wee1-HA were incubated with 100 ng/ml LMB for 1 h. Total protein lysates were resolved by SDS-PAGE and membranes probed with the indicated antibodies. **G.** Cells expressing Mik1-HA were exposed to 10 mM caffeine and harvested at the indicated time points. Samples were treated as in **A**.

### Caffeine suppresses DNA damage-induced Wee1 accumulation

Exposure to DNA damaging agents has previously been shown to induce the accumulation of Wee1 (Raleigh & O’Connell, 2000). We thus studied the effect of caffeine on Wee1 expression under these conditions. Incubation with 10 mM caffeine alone induced the accumulation of Wee1 but had no impact on the slight suppressive effect of 20 mM HU on the protein (Fig. 3 A). Exposure to 10 µg/ml phleomycin induced Wee1 accumulation in a manner akin to that of caffeine. Interestingly, caffeine strongly inhibited phleomycin-induced Wee1 accumulation such that the levels were even below those observed in untreated cells (Fig. 3 A). Caffeine activates both the DNA damage response and environmental stress response pathways (Calvo *et al*., 2009, Alao *et al*., 2014). The effect of caffeine on Wee1 stability may thus be context dependent. The loss of Wee1 expression in the presence of phleomycin would be expected to override checkpoint signalling under these conditions. Caffeine-induced Wee1 accumulation was dependent on Rad24 expression (Fig. 3 B). Exposure to HU in *rad24*Δ mutants did not induce Wee1 accumulation, and co-exposure to caffeine completely abolished expression of this kinase. Wee1 accumulation in response to phleomycin exposure was not observed in *rad24*Δ mutants. In marked contrast to wt cells, caffeine had no effect on Wee1 expression in *rad24*Δ mutants exposed to phleomycin (Fig. 3 B). In fact, the phleomycin-induced accumulation of Wee1 did not occur in *rad24*Δ mutants, identifying a direct role for Rad24 in regulating DNA damage checkpoints (Fig. 3 B). We next compared the effect of phleomycin exposure on Cdc2 Tyr 15 phosphorylation in wt and *rad24*Δ mutants. Exposure to 10 µg/ml phleomycin increased the basal level of Cdc2 phosphorylation (Fig. 3 C). In *rad24*Δ mutants, basal Cdc2 phosphorylation was not detected and exposure to phleomycin resulted in levels of Cdc2 phosphorylation below those of untreated wt cells (Fig. 3 C). We previously demonstrated that caffeine stabilises Cdc25 in the nucleus of S. pombe cells exposed to genotoxic agents (Alao *et al*., 2014). Caffeine thus both stabilises Cdc25 and suppresses Wee1 expression under genotoxic conditions. This activity would effectively lead to the abolition of the “double lock” DNA damage checkpoint mechanism.

**Figure 3.**
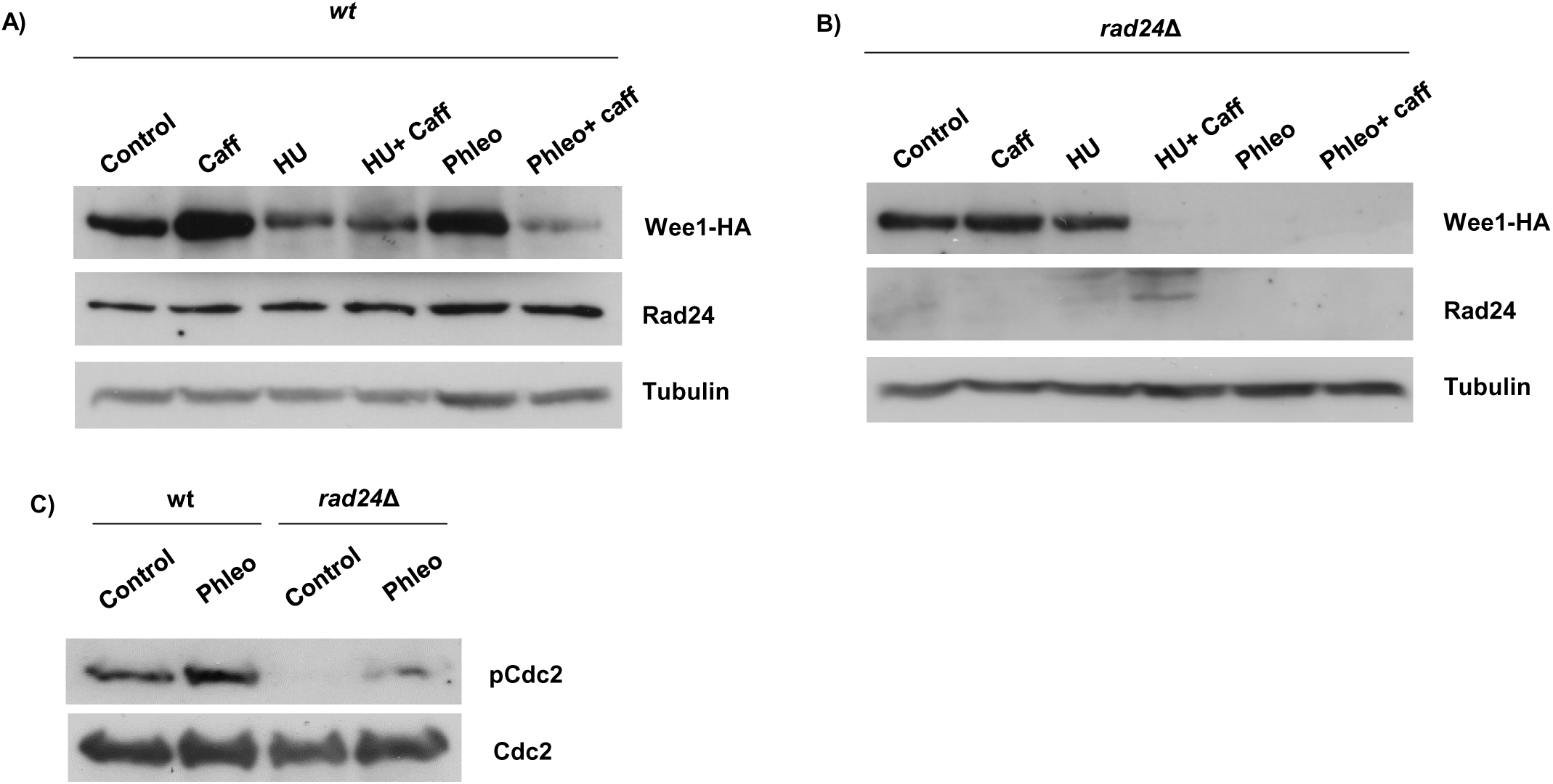
Caffeine suppresses genotoxin-induced Wee1 accumulation in *S. pombe*. **A.** Cells expressing Wee1-HA were cultured with 10 mM caffeine alone or in combination with 20 mM HU or 10 µg/ml phleomycin as indicated. Cultures were pre-treated with HU or phleomycin for 2 h and then for a further 2 h in the presence or absence of caffeine. **B.** Wee1-HA *rad24*Δ cells were treated as in A. **C.** Wt cells expressing Wee1-HA and Wee1-HA *rad24*Δ cells were exposed to 10 µg/ml phleomycin for 1 h. Total protein lysates were resolved by SDS-PAGE and membranes probed with antibodies against phospho-and total Cdc2.

### Caffeine mediates checkpoint override by suppressing Wee1 under genotoxic conditions

We previously observed that the effect of caffeine on cell cycle progression in *S. pombe* is enhanced in *wee1*Δ and other checkpoint mutants (Alao *et al*., 2014). Caffeine (10 mM) overrode checkpoint signalling in *wee1*Δ mutants but to a lesser extent than in wt cells (Fig. 4 A). As judged by FACS analyses, survival assays, and cell morphology, caffeine only slightly increases the sensitivity of *wee1*Δ mutants to 20 mM HU relative to wt cells (Fig. 4 B-D). We did not detect a differential level of sensitivity to HU in *rad24*Δ and *wee1*Δ mutants relative to wt cells (Fig. 4 E). Unlike wt cells and *wee1*Δ mutants however, *rad24*Δ mutants did not become elongated following exposure to HU (Fig. 4 E). Interestingly, caffeine was far more effective at overriding checkpoint signalling in response to HU in *rad24*Δ mutants (Fig. 4 G,I). We also observed that in contrast to HU, *rad24*Δ and *wee1*Δ mutants are highly sensitive to phleomycin (Fig. 4 H).

**Figure 4.**
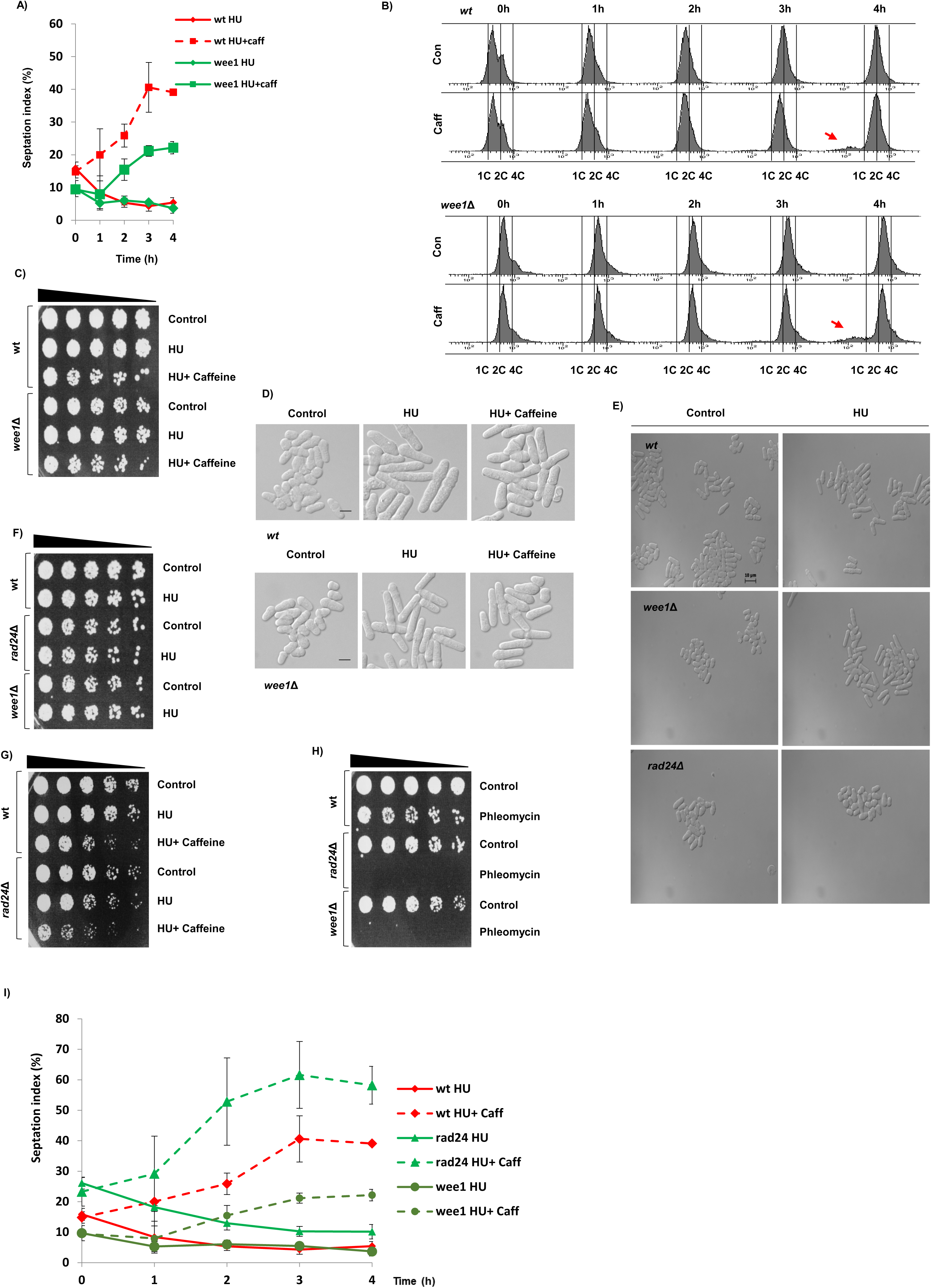
Differential effects of caffeine on cell cycle progression in *S. pombe wee1*Δ and *rad24*Δ mutants. **A.** Duplicate cultures of wt and *wee1*Δ cells were incubated with 10 µg/ml phleomycin for 2 h. The cultures were then incubated for a further 4 h with or without 10 mM caffeine and samples harvested at the indicated time points. Cells were fixed in 70 % ethanol, stained with aniline blue and the septation index determined by fluorescent microscopy. **B.** Samples from A were stained with propidium iodide and analysed by FACS. Arrows indicate cell with mis-segregated chromosomes. **C.** Wt and *wee1*Δ cells were incubated with 20 mM HU for 2 h. The cultures were then incubated for a further 2 h in the presence of 10 mM caffeine as indicated. Cultures were adjusted for relative cell numbers, serially diluted and plated unto YES agar plates. Plates were incubated at 30°C for 2 - 3 days. **D.** Wt and *wee1*Δ cells were treated as in A. Cells were fixed in 70 % ethanol and examined by differential contrast microscopy. **E.** Wt and *rad24*Δ *wee1*Δ cells were exposed to 20 mM HU for 4 h. Cells were fixed in 70 % ethanol and examined by differential contrast microscopy. **F,G**. Wt and *rad24*Δ *wee1*Δ cells were treated as in **A**. **H**. Wt and *rad24*Δ *wee1*Δ cells were exposed to 10 µg/ml phleomycin for 2 h. Cultures were adjusted for relative cell numbers, serially diluted and plated unto YES agar plates. Plates were incubated at 30° C for 2 - 3 days.

### Inhibition of TORC1 signalling overrides checkpoint signalling

The TORC1 complex regulates the timing of mitosis and its inhibition by rapamycin or torin1 leads to an advanced entry into mitosis (Petersen & Nurse, 2007, Atkin *et al*., 2014). As caffeine inhibits TORC1 and advances entry into mitosis (Rallis *et al*., 2013, Alao *et al*., 2014), we investigated if TORC1 inhibition similarly overrides checkpoint signalling. Exposure to 20 mM HU or 7.5 µM torin1 for 4 h did not affect the viability of wt *S. pombe* cells (Fig. 5 A). Unlike caffeine, co-exposure with torin1 did not affect the sensitivity of wt cells to HU (Figs. 4 G and 5 A). Cells co-exposed to HU and torin1 were shorter than cells exposed to HU alone but no chromosome mis-segregation was observed. In contrast, cells exposed to torin1 alone displayed a “wee” phenotype (Fig. 5 B).

**Figure 5.**
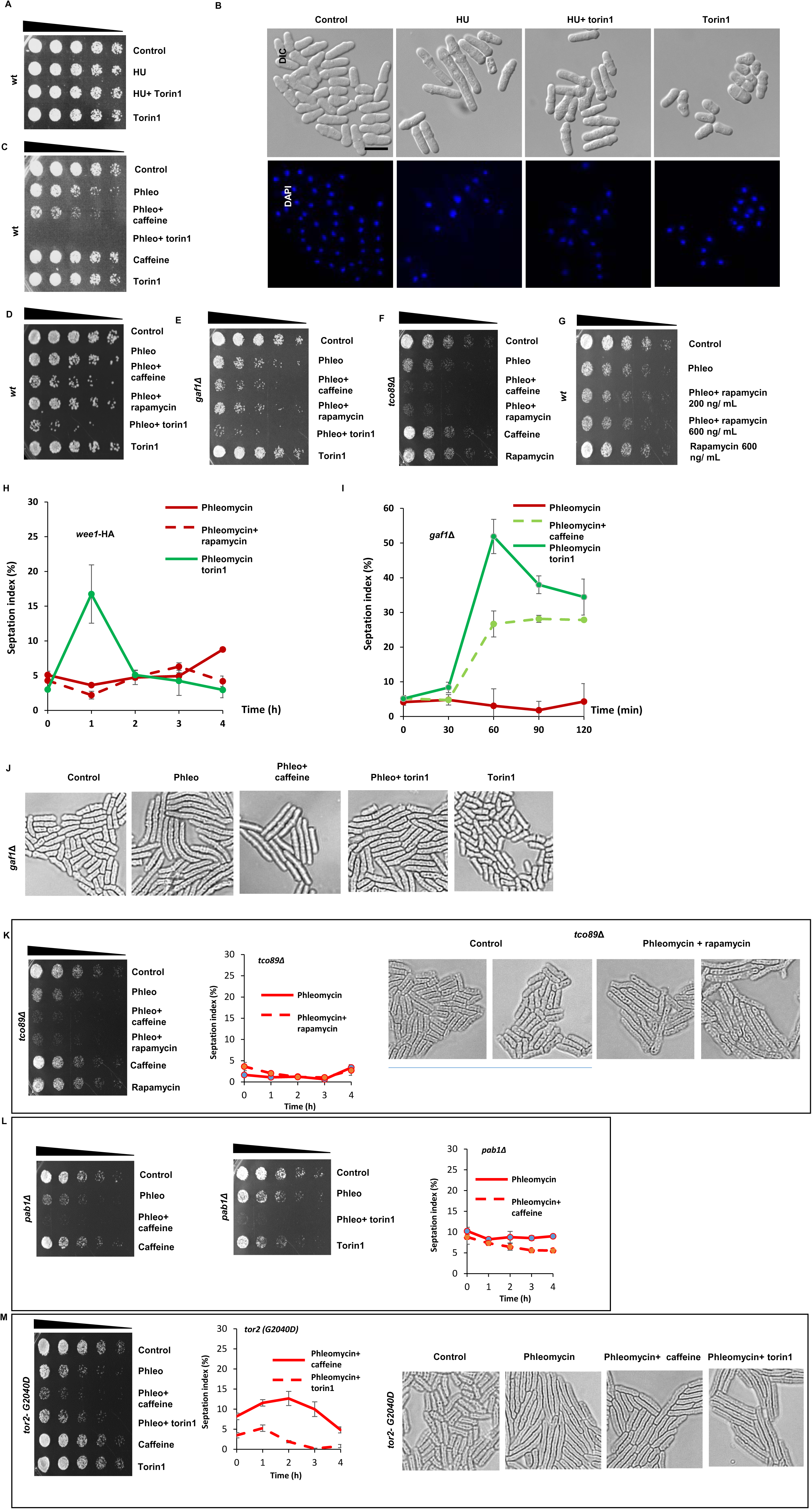
Inhibition of TORC1 overrides checkpoint signalling similarly to caffeine. **A.** Wt cells were incubated with 20 mM HU for 2 h. The cultures were then incubated for a further 2 h in the presence of 10 mM caffeine as indicated. Cultures were adjusted for relative cell numbers, serially diluted and plated unto YES agar plates. Plates were incubated at 30° C for 2 - 3 days. **B.** Cells in A were fixed in 70 % ethanol, stained with DAPI and examined by differential contrast microscopy. **C.** Wt cells were incubated with 10 µg/m phleomycin for 2 h. The cultures were then incubated for a further 2 h in the presence of 10 mM caffeine or 7.5 µM torin1 as indicated. Cultures were adjusted for relative cell numbers, serially diluted and plated on YES agar plates. Plates were incubated at 30° C for 2- 3 days. **D.** Wt cells expressing Wee1-HA, were incubated with 5 µg/ml of phleomycin for 2 h. The cultures were then incubated for a further 2 h in the presence of 10 mM caffeine, 200 ng/ml rapamycin or 5 µM torin1 as indicated. Cultures were adjusted for relative cell numbers, serially diluted and plated on YES agar plates. Plates were incubated at 30° C for 2 - 3 days. **E.** *gaf1*Δ mutant cells were incubated with 5 µg/ml of phleomycin for 2 h. The cultures were then incubated for a further 2 h in the presence of 10 mM caffeine, 200 ng/ml rapamycin or 5 µM torin1 as indicated. Cultures were adjusted for relative cell numbers, serially diluted and plated on YES agar plates. Plates were incubated at 30°C for 2 - 3 days. **F.** *tco89*Δ mutant cells were incubated with 5 µg/ml of phleomycin for 2 h. The cultures were then incubated for a further 2 h in the presence of 10 mM caffeine, 200 ng/ml rapamycin or 5 µM torin1 as indicated. Cultures were adjusted for relative cell numbers, serially diluted and plated onto YES agar plates. Plates were incubated at 30°C for 2 - 3 days. **G.** Wt cells were exposed to 5 µg/ml of phleomycin for 2 h. The cultures were then incubated for a further 2 h in the presence of 200 ng/ml or 600 ng/ml rapamycin as indicated. Cells exposed to 600 ng/ml rapamycin served as a control. Cultures were adjusted for relative cell numbers, serially diluted and plated on YES agar plates. Plates were incubated at 30° C for 2 - 3 days. **H.** Wt cells were incubated with 5 µg/ml of phleomycin for 2 h. The cultures were then incubated for a further 2 h in the presence of 200 ng/ml rapamycin or 5 µM torin1. Samples were harvested at the indicated time points. Cells were fixed in 70 % ethanol, stained with calcofluor white and the septation index determined by fluorescent microscopy. **I.** *gaf1*Δ mutants were incubated with 5 µg/ml of phleomycin for 2 h. The cultures were then incubated for a further 2 h in the presence of 10 mM caffeine or 5 µM torin1. Cells were fixed in 70 % ethanol, stained with aniline blue or calcofluor white and the septation index determined by fluorescent microscopy. **J.** *gaf1*Δ mutants were incubated with 5 µg/ml of phleomycin for 2 h. The cultures were then incubated for a further 2 h in the presence of 10 mM caffeine or 5 µM torin1. Cells were fixed in 70 % ethanol, stained with DAPI and examined by differential contrast microscopy. **K.** *tco89*Δ mutants were incubated with 5 µg/ml of phleomycin for 2 h. The cultures were then incubated for a further 2 h in the presence of 10 mM caffeine or 200 ng/ml rapamycin. Cultures were adjusted for relative cell numbers, serially diluted and plated on YES agar plates. Plates were incubated at 30° C for 2 - 3 days. Alternatively, cells were treated with 10 mM caffeine or 200 ng/ mL of rapamycin alone. Samples were harvested at the indicated time points. Cells were fixed in 70 % ethanol, stained with aniline blue or calcofluor white and the septation index determined by fluorescent or the relative cell lengths with differential contrast microscopy. **L.** *pab1*Δ mutants were incubated with 5 µg/ml of phleomycin for 2 h. The cultures were then incubated for a further 2 h in the presence of 10 mM caffeine or 5 µM torin1. Alternatively, cells were treated with 10 mM caffeine or 5 µM torin1 alone. Cultures were adjusted for relative cell numbers, serially diluted and plated on YES agar plates. Plates were incubated at 30° C for 2 - 3 days. Alternatively, samples were harvested at the indicated time points. Cells were fixed in 70 % ethanol, stained with aniline blue or calcofluor white and the septation index determined by fluorescent microscopy. **M.** The *tor2*(*G2040D*) strain was incubated with 5 µg/ml of phleomycin for 2 h. The cultures were then incubated for a further 2 h in the presence of 10 mM caffeine or 5 µM torin1. Cultures were adjusted for relative cell numbers, serially diluted and plated on YES agar plates. Plates were incubated at 30° C for 2 - 3 days. Alternatively, samples were harvested at the indicated time points. Cells were fixed in 70 % ethanol, stained with aniline blue or calcofluor white and the septation index determined by fluorescent or the relative cell lengths with differential contrast microscopy.

In contrast to its effects on HU sensitivity, torin1 was far more effective at sensitising wt cells to phleomycin than caffeine (Fig. 5 C-D). Similar results were obtained with *gaf1*Δ mutants, a transcription factor that partially mediates the effect of torin1 (Laor *et al*., 2015, Rodríguez-López *et al*., 2019), (Fig. 5 E,I). In addition, *gaf1*Δ mutants exposed to phleomycin and torin1 were shorter than cell exposed to phleomycin alone (Fig. 5 J). Torin1 thus overrides DNA damage checkpoint signalling independently of Gaf1. Rapamycin failed to override checkpoint signalling concentrations as high as 600 ng/ml in wt *S. pombe* cells (Fig. 5 G). Tco89 is a subunit of the TORC1 complex and *tco89*Δ mutants are hypersensitive to caffeine and rapamycin (Reinke *et al*., 2006, Hayashi *et al*., 2007). Rapamycin sensitised *tco89*Δ mutants to phleomycin in a manner similar to caffeine (Fig. 5 F). Further studies demonstrated, however, that rapamycin does not enhance sensitivity to phleomycin by overriding DNA damage checkpoint signalling (Fig. 5). Active TORC1 delays the timing of mitosis, by indirectly inhibiting the PP2A phosphatase catalytic subunit Pab1. Thus, TORC1 inhibition results in advanced mitosis and a smaller cell size (Martin & Lopez-Aviles, 2018). Caffeine and torin1 were still able to enhance sensitivity of *pab1*Δ mutants to phleomycin (Fig. 5 L). As with *tco89*Δ mutants, this enhanced sensitivity to phleomycin did not result in DNA damage checkpoint override (Fig. 5 L). To confirm that torin1 overrides checkpoint signalling in a TORC1 resistant manner, we investigated its effect on the torin1-resistant *tor2-G2040D* mutant (Atkin *et al*., 2014). Unlike caffeine, torin1 failed to override phleomycin-induced DNA damage checkpoint signalling in the *tor2-G2040D* mutant (Fig. 5 M). Together with previous findings (Reinke *et al*., 2006, Wanke *et al*., 2008, Alao *et al*., 2014), our study strongly suggests that caffeine overrides DNA damage checkpoint signalling by targeting Tor2 and the TORC1 complex. We cannot however, rule out that caffeine can enhance sensitivity to DNA damage independently of checkpoint override, as DNA repair pathways are required for resistance to the drug (Calvo *et al*., 2009).

## Discussion

The precise mechanisms whereby caffeine overrides DNA damage checkpoint signalling remain unclear. In the present study, we investigated further how caffeine modulates cell cycle progression through Cdc25 and Wee1. Initial studies suggested that caffeine inhibits Rad3 and its related homologues (Moser *et al*., 2000, Jimenez *et al*., 1992). Caffeine however exerts inhibitory activity on several members of the phosphatidylinositol 3 kinase-like kinase (PIKK) family (Humphrey, 2000, Bode & Dong, 2007, Lovejoy & Cortez, 2009, Gibbs *et al*., 2015)). More recent studies suggest that the Tor2-containing TORC1 complex is the major target of caffeine *in vivo* (Kuranda *et al*., 2006, Reinke *et al*., 2006, Wanke *et al*., 2008, Rallis *et al*., 2013). TORC1 regulates the timing of mitosis by modulating Cdc25 and Wee1 activity. Inhibition of TORC1 activity thus advances cells into mitosis, an effect like that observed with caffeine (Atkin *et al*., 2014, Alao *et al*., 2014). DNA damage checkpoint activation and enforcement require the dual inhibition of Cdc25 activity and activation of Wee1 (Raleigh & O’Connell, 2000). We previously demonstrated that caffeine induces Cdc25 accumulation independently of Rad3 inhibition (Alao *et al*., 2014). As TORC1 regulates Cdc25 and Wee1 activity and is inhibited by caffeine, this inhibition may in fact underlie the effects of the compound on cell cycle progression.

### Effect of caffeine on Cdc25 stability

Cdc25 undergoes Cdc2-dependent activating phosphorylation as well as inhibitory phosphorylation via the Rad3 and Sty1 regulated signalling pathways (Perry & Kornbluth, 2007). Caffeine stabilises wt and Cdc25 mutant proteins that cannot be phosphorylated in response to DNA damage. Caffeine thus clearly stabilises Cdc25 independently of its negative phosphorylation. Furthermore, caffeine is more effective at stabilising the Cdc25_(9A)_- GFP_int_ and Cdc25_(12A)_-GFP_int_ isoforms and hence checkpoint override in these genetic backgrounds. The stabilising effect of caffeine on Cdc25 is also independent of Sty1 signalling, as exposure to osmotic stress induced the degradation of Cdc25_(9A)_-GFP_int_. The Cdc25_(9A)_-GFP_int_ and Cdc25_(12A)_-GFP_int_ mutants are also rapidly degraded following exposure to genotoxic agents (Frazer & Young, 2011, Frazer & Young, 2012, Alao *et al*., 2014). These conditions must thus cause cellular changes that result in the targeting of these mutants for ubiquitin-dependent degradation. Inhibition of nuclear export with LMB resulted in a decrease in Cdc25 levels and prevented its stockpiling following exposure to HU. LMB also failed to prevent the HU-induced degradation of the Cdc25_(12A)_-GFP_int_ mutant protein. As expected, caffeine-induced Cdc25 stabilisation was independent of Rad24. Caffeine thus appears to partially inhibit the nuclear degradation of Cdc25 in *S. pombe*. Accordingly, Cdc25_(12A)_-GFP_int_ mutants are more susceptible to caffeine-mediated checkpoint override than wt cells. It remains unclear if caffeine-induced Cdc25 accumulation results from TORC1 inhibition.

### Effect of caffeine on Wee1 stability

Our previous studies suggested that Wee1 attenuates the effect of caffeine on cell cycle progression in *S. pombe* (Alao *et al*., 2014). Furthermore, Cdc25 and Wee1 are co-regulated during the cell cycle and thus determine the timing of mitosis (Atkin *et al*., 2014, Lucena *et al*., 2017). We thus investigated the effect of caffeine on Wee1 expression. Interestingly, caffeine induced rapid Wee1 accumulation under normal growth conditions. This accumulation was dependent on Rad24 expression which was also induced by exposure to caffeine (Fig. 2 A-C). Sty1 was recently shown to modulate the ratio of Cdc25 to Wee1 in a Rad24-dependent manner (Paul *et al*., 2018). Deletion of *rad24^+^* resulted in reduced Wee1 expression indicating that unlike Cdc25, Wee1 stability is dependent on Rad24 under normal cell cycle conditions (Figs. 1 E, 2 D and refs. (Frazer & Young, 2011, Frazer & Young, 2012, Alao *et al*., 2014)). Deletion of *rad24^+^* resulted in a “semi-wee” phenotype. These findings suggest that it is the lack of Wee1 expression rather than constitutively nuclear Cdc25 expression, that is responsible for the shorter length at division observed in *rad24*Δ mutants (Fig. 2 D and ref. (Ford *et al*., 1994)). Inhibiting nuclear export with LMB also stabilised Wee1 independently of Rad24. Further, caffeine stabilised the Mik1 kinase, an S-phase specific inhibitor of Cdc2 (Lundgren *et al*., 1991). Caffeine thus appears to stabilise Cdc25, Mik1 and Wee1 under normal cell cycle conditions. We previously demonstrated that the effect of caffeine on cell cycle progression is dampened by Srk1 and Wee1 activity (Alao *et al*., 2014). It remains unclear if Rad24 stabilises Mik1 and how caffeine affects this interaction in the presence of stalled replication forks or DNA damage. Remarkably, the effect of caffeine on Cdc25 and Wee1 is reversed under genotoxic conditions. Under these conditions Cdc25 is normally inactivated and sequestered in the cytoplasm or degraded in the nucleus, while Wee1 becomes activated and accumulates in the nucleus (Frazer & Young, 2011, Frazer & Young, 2012). Exposure to caffeine under these conditions results in the stabilisation of Cdc25 within the nucleus (Alao *et al*., 2014) and degradation of Wee1 (Fig. 3 A,B). Exposure to phleomycin but not HU induced Wee1 accumulation in a Rad24-dependent manner. In contrast to caffeine or phleomycin exposure alone, Wee1 did not accumulate when the cells were co-exposed to both compounds. Interestingly, caffeine also abolished Wee1 expression in *rad24*Δ mutants exposed to HU. Total phospho-Cdc2 levels are also suppressed in *rad24*Δ mutants, consistent with a loss of Wee1 expression and a “semi-wee” phenotype. The TORC1 complex also regulates the activity of Cdc25 and Wee1 under normal cell cycle conditions to regulate the timing of mitosis (Atkin *et al*., 2014). Furthermore, Sty1 can modulate the relative expression levels of Cdc25 and Wee1 in a Rad24-dependent manner (Paul *et al*., 2018). Crosstalk between TORC1, Sty1, and the replication checkpoint pathway has also been reported (Hartmuth & Petersen, 2009, Fletcher *et al*., 2018). As caffeine inhibits TORC1 and activates Sty1, its effect on cell cycle progression may be context dependent and result from fundamental changes to physiological co-regulation of Cdc25 and Wee1. Caffeine-mediated TORC1 inhibition may also influence autophagy and 26S proteasomal degradation (Gressner, 2009, Marshall & Vierstra, 2015, Zhao *et al*., 2015). In any case, caffeine clearly abolishes Cdc25 inhibition and degradation under genotoxic conditions independently of Rad24. In contrast, Wee1 degradation may result from changes to its phosphorylation and ability to interact with Rad24. Caffeine thus overrides the DNA damage “double lock” mechanism independently of Rad3 inhibition ((Alao *et al*., 2014) and this study).

### Differential effects of caffeine on DNA damage resistance

The sensitivity of *rad24*Δ and *wee1*Δ mutants following a 4-hour exposure to HU was not enhanced relative to wt cells (Fig. 4 F). Caffeine was however no more effective at driving checkpoint override in *wee1*Δ mutants exposed to HU than in wt cells. This may reflect the differential cell cycle kinetics of *wee1*Δ mutants which delay progression through G1, because of size constrains and the more important role of Mik1 under these conditions (Frazer & Young, 2011, Frazer & Young, 2012). Future studies will investigate the effect of caffeine on Mik1 expression in cells exposed to HU. Alternatively, the increase in Cdc25 activity induced by caffeine in a *wee1*Δ background may delay progression through cytokinesis due to high Cdc2 activity. Caffeine was more effective at overriding the replication checkpoint in *rad24*Δ mutants compared to wt cells. Rad24 binding and nuclear export are not required for the inhibition of Cdc25 activity. Caffeine overrides checkpoints more efficiently in mutants expressing Cdc25 isoforms that cannot be phosphorylated (Alao *et al*., 2014). It remains unclear if Rad24 stabilises Mik1 in a manner like Wee1. Increased nuclear levels of Cdc25, following exposure to HU combined with decreased Mik1 expression, might account for the greater effect of caffeine on *rad24*Δ mutants. Caffeine may also suppress Mik1 expression similarly to Wee1 (unpublished results). The sensitivity of *rad24*Δ and *wee1*Δ mutants to phleomycin was identical, probably reflecting the lack of Wee1 expression in these genetic backgrounds.

### TORC1 inhibition overrides DNA damage checkpoint signalling

Recent studies have suggested that TORC1 and not Rad3 and its homologues is the preferred target of caffeine in vitro (Reinke *et al*., 2006, Wanke *et al*., 2008, Alao *et al*., 2014). TORC1 regulates the timing of mitosis by regulating the activity of the PP2A phosphatase, which in turn regulates the activity of Cdc25 and Wee1 (Atkin *et al*., 2014, Martin & Lopez-Aviles, 2018). Exposure of *S. pombe* cells to rapamycin or torin1, activates Cdc25 and suppresses the expression of Wee1, resulting in advanced entry into mitosis. Furthermore, the effect of caffeine on cell cycle progression mimics that of rapamycin and torin1 and is dependent of Cdc25 (Alao *et al*., 2014, Atkin *et al*., 2014). We thus hypothesised that TORC1 inhibition by rapamycin or torin1 should override DNA damage checkpoint signalling in a manner akin to caffeine. In wt, caffeine, rapamycin and torin1 all advance the timing of mitosis under normal cell cycle conditions in *S. pombe* (Alao *et al*., 2014, Atkin *et al*., 2014). In this study, caffeine and torin1 but not rapamycin overrode phleomycin-induced DNA damage checkpoint activation. Interestingly torin1 did not enhance sensitivity to HU in this context. This may be due to differential effects of caffeine on additional signalling pathways (*e.g.* Mik1 expression and global 26S proteasome-mediated protein degradation (Alao et al., unpublished results). The TORC1 downstream transcription factor Gaf1 has recently been shown to mediate the effects of Tor2 inhibition on chronological lifespan in *S. pombe* (Laor *et al*., 2015, Rodríguez-López *et al*., 2019). Caffeine and torin1 clearly increased sensitivity to phleomycin in *gaf1*Δ mutants, suggesting this effect occurs independently of Gaf1. Rapamycin could enhance sensitivity to the *tco89*Δ mutant which displays enhanced sensitivity to the drug and other genotoxins (Nakashima *et al*., 2012, Pan *et al*., 2012). This activity was not associated with DNA damage checkpoint override. TORC1 inhibition results in advanced entry into mitosis via indirect inhibition of the PP2A phosphatase and its Pab1 activating subunit (Atkin *et al*., 2014, Martin & Lopez-Aviles, 2018). Caffeine and torin1 enhanced DNA sensitivity in a *pab1*Δ genetic background, albeit independently of checkpoint override. These observations suggest that caffeine and torin1 induce checkpoint override in wt but not *pab1*Δ mutants. The increased sensitivity to DNA damage in these genetic backgrounds suggests that caffeine, and by implication torin1, can enhance sensitivity to genotoxins independently of checkpoint override. Indeed, resistance to caffeine requires Rad3 and downstream modulators of DNA damage repair such as Rhp51 and Rhp54 (Calvo *et al*., 2009). Our finding, that caffeine but not torin1 overrides checkpoint signalling in the torin1 resistant *tor2-G2040D* mutant (Atkin *et al*., 2014) confirms that TORC1 inhibition overrides checkpoint signalling in the presence of genotoxins.

Effective G2 checkpoint activation requires the dual inhibition of Cdc25 and activation of Wee1 (Raleigh & O’Connell, 2000). Mutants that fail to express *wee1* and *rad24* are especially sensitive to DNA damage as these genes regulate the G2 checkpoint. Both rapamycin and torin1 suppress Wee1 expression in *S. pombe*, although rapamycin is less effective in this regard (Atkin *et al*., 2014). We have demonstrated that caffeine induces Wee1 expression under normal cell cycle conditions in a Rad24-dependent manner. Curiously, this effect is reversed under genotoxic conditions, where co-exposure to caffeine prevents Wee1 accumulation. The failure of rapamycin to override checkpoint signalling may thus result from its less effective inhibition of TORC1 and Wee1 suppression relative to torin1 and caffeine. Indeed, rapamycin effectively overrode DNA damage checkpoint signalling in *tco89*Δ mutants that display hypersensitivity to the inhibitor (Reinke *et al*., 2006, Hayashi *et al*., 2007). Taken together our finding and those of others, strongly suggest that caffeine overrides DNA damage signalling independently of Rad3 by inhibiting TORC1 activity.

## Experimental procedures

### Strains, media and reagents

Strains are listed in Table 1. Cells were grown in yeast extract plus supplements medium (YES) Stock solutions of caffeine (Sigma Aldrich AB, Stockholm, Sweden) (100 mM) were prepared in water stored at −20°C. HU (Sigma Aldrich AB) was dissolved in water at a concentration of 1 M and stored at −20°C. Phleomycin (Sigma Aldrich AB) was dissolved in water and stock solutions (10 µg/ml) stored at −20°C. Alternatively, phleomycin was purchased from Fisher scientific UK as a 20 mg/ mL solution and stored at −20°C.

**Table 1.** S. pombe strains

|  |  |
| --- | --- |
| *h<sup>-</sup>* L972 | Lab stock |
| *h<sup>+</sup>* *cdc25-6HA [ura4<sup>+</sup>] leu1-32 ura4-D18* (FY7031) | YGRC |
| *h<sup>+</sup>* *cdc25-6HA [ura4<sup>+</sup>] leu1-32 ura4-D18 rad24::KanMX6* | This study |
| *h<sup>-</sup>* *cdc25-12myc::ura4<sup>+</sup> ura4-D18 leu1-32* | P. Russell |
| *h<sup>-</sup>* *cdc25-GFP<sub>int</sub> cdc25::ura4<sup>+</sup> ura4-D18 leu1-32* | P. Young |
| *h<sup>+</sup>* *cdc25<sub>(9A)</sub>-GFP<sub>int</sub> cdc25::ura4<sup>+</sup> ura4-D18 leu1-32* | P. Young |
| *h<sup>-</sup>* *cdc25<sub>(12A)</sub>-GFP<sub>int</sub> cdc25::ura4<sup>+</sup> ura4-D18 leu1-32* | P. Young |
| *h<sup>-</sup>* *cdc25<sub>(12A)</sub>-GFP<sub>int</sub> cdc25::ura4<sup>+</sup> ura4-D18 leu1-32 mik1::ura4<sup>+</sup>* | P. Young |
| *h<sup>+</sup>* *cut8::ura4* (FY9535) | YGRC |
| *h<sup>+</sup>* *leu1 his2 ura4 cut8-8xMyc ura4<sup>+</sup>* | YGRC |
| *h<sup>-</sup>* *wee1::ura4<sup>+</sup> leu1-32 ura4-D18* (FY7283) | YGRC |
| *h<sup>-</sup>* *wee1-3HA:6His leu1-32 ura4-D18* (FY16241) | YGRC |
| *h<sup>-</sup>* *wee1-3HA:6His leu1-32 ura4-D18 rad24::KanMX6* | This study |
| *h<sup>-</sup>* *rad24::ura4<sup>+</sup> leu1 ura4-D18 ade6-M210* (FY13517) | YGRC |
| *h<sup>-</sup>* *mik1::ura4 leu1 ura4* (FY8317) | YGRC |
| *h<sup>+</sup>* *ade6-M210 ura4-D18 leu1-32 gaf1::KanMX6* | Bioneer |
| *h<sup>+</sup>* *ade6-M210 ura4-D18 leu1-32 tco89::KanMX6* | Bioneer |
| *h<sup>+</sup>* *leu1-32 ura4-D18Δ pab1::kanR* | D. R. Kellogg |
| *h<sup>+</sup>* *tor2-G2040D* | J. Petersen |

YGRC, Yeast Genetic Resource Center, Osaka, Japan

### Molecular genetics

Deletion of the open reading frames was done by PCR-based genomic targeting using a *KanMX6* construct (Bähler *et al*., 1998). Disruptions were verified by PCR using genomic DNA extracted from mutants.

### Microscopy

Calcofluor white (Sigma-Aldrich) staining and septation index assays were carried out as previously described (Alao *et al*., 2014, Dunaway & Walworth, 2004, Forsburg & Rhind, 2006). Images were obtained with a Zeiss AxioCam on a Zeiss Axioplan 2 microscope with a 100 × objective using a 4,6-diamidino-2-phenylindole (DAPI) filter set.

### Fluorescence-activated cell sorting (FACS)

Cells were harvested at the desired time points, resuspended in 70 % ethanol and stored at 4°C until use. FACS analyses were performed according to the previously described protocol (Alao *et al*., 2014), using propidium iodide (32 µg/ml) as outlined on the Forsburg lab page (http://www-rcf.usc.edu/~forsburg/yeast-flow-protocol.html). Flow cytometry was performed with a BD FACSAria™ cell sorting system (Becton Dickinson AB, Stockholm, Sweden).

### Immunoblotting

Monoclonal antibodies directed against HA (F-7), Myc (9E10) and pan 14-3-3 (K-19) proteins were from Santa Cruz Biotechnology (Heidelberg, Germany). Monoclonal antibodies directed against GFP (11814460001) and α-tubulin were from Sigma-Aldrich (Sigma Aldrich AB). Polyclonal antibodies directed against phospho-(Tyr15) Cdc2 were from Cell Signaling Technology (BioNordika, Stockholm, Sweden). Monoclonal antibodies against Cdc2 were from Abcam (Cambridge, UK). For immunoblotting, protein extracts were prepared as previously described (Alao *et al*., 2014) with addition of 1 × PhosStop phosphatase inhibitor cocktail (Roche Diagnostics Scandinavia AB, Bromma, Sweden). Proteins were separated by SDS-PAGE. Epitope-tagged proteins were detected with the appropriate monoclonal antibodies.

## Author’s contributions

J.P.A., C.R. and P.S. conceived and designed the study. J.P.A. and J.J.S. carried out experiments and analyzed the data. J.P.A. wrote the manuscript. All authors read, revised and approved the final manuscript.

## Acknowledgements

We are grateful to S. Ali, R. Lucena, D. R. Kellogg and J. Petersen for technical support. This work was financially supported by Carl Trygger’s Foundation (CTS 15:13) and the Swedish Cancer Fund (13-0438 and 16-0708). CR is funded by UEL QR funds.

